# Diet and Domestication Drive Evolution of the Gut Holobiome

**DOI:** 10.1101/2021.10.14.464420

**Authors:** Vivek Ramanan, Shanti Mechery, Indra Neil Sarkar

**Affiliations:** Center for Biomedical Informatics, Brown University, Providence, Rhode Island, USA; Rhode Island Quality Institute, Providence, Rhode Island, USA

## Abstract

The host microbiome encompasses all microorganisms of a host. Host and microbiome coevolution in the gut result in differing microbial compositions, functionality, and host diet [1]. Host diet modulates what macromolecules are used for gut microbial metabolism, which can determine digestion, health, and behavior [2, 3]. Microbial composition across animals provides data on how microbiomes segregate between species and diets [4]. Here we show that microbiome data from GenBank can model host evolution, providing a “holobiome” insight to the important roles of diet and domestication. The main findings of this study in respect to microbial composition among species were: (1) herbivores are more similar than hosts with other diets; (2) domesticated species are more similar than wild relatives; and (3) humans are distinct from primates. Microbial composition between diets indicates a difference in functionality, where protein and fiber degradation are seen more in carnivores and herbivores respectively. Additionally, herbivores show the most microbial diversity among the diets. Finally, this analysis informs us of gaps in current microbiome data collection, which is biased toward pathogens. Thus, the host-microbiome relationship depicts a complex web of microbial functionality, composition, and diet that impact coevolution.

## Introduction

The microbiome plays a significant role in shaping hosts on an individual and species scale, demonstrating how complex evolutionary forces inform the coevolution of humans with their microbiomes. Prior studies have explored microbiome-host coevolution, including those examining primate evolution. The combined evolutionary concept of host species and coevolution with their symbionts is referred to as the “holobiome” [5]. The holobiome can be influential at multiple evolutionary levels, and used to examine microbiome-host coevolution for identifying potential influential factors using phylogeny [6]. The holobiome presumes that microbial symbionts and host fitness are interdependent, therefore enabling phylosymbiosis to occur, where the evolution of one parallels the evolution of the other [1, 7, 8]. Phylosymbiotic principles suggest that the microbiome can be the basis for holobiome phylogenetic tree construction to identify evolutionarily linked features for a set of host species.

Gut microbiome composition across host evolution portrays how bacterial lineages impact host diet [9]. Brooks, *et al*. found that within-species microbiome variation is less than between-species microbiome variation. They further found that within-species microbiota transplants can impact survival and performance reductions depending on the relatedness of the host-species pairs [10]. Certain microbial clades are consistent across mammals, where bacterial lineages are aligned with host phylogeny [11]. Microbiome analysis has been used to examine potential paths for bacterial influence in the divergence of humans from other primates. In characterizing the microbiome impact on human evolution, Davenport *et al*. analyzed the human microbiome alongside old world and new world primates [4]. These comparisons revealed that the human microbiome is closer to old world primates than new world primates. Subsequent studies have shown how human microbiomes diverged from primate microbiomes, likely due to a loss of microbial diversity and diet changes [12]. Analysis of mammalian microbiomes has shown that microbiome composition can be linked to diet, which also indicates that bacterial diversity is greatest in herbivores who consume more dietary fiber [13]. This research demonstrates the importance of microbiome analysis for evolutionary history and that host evolution is tied to microbiome lineages, diet, and diversity.

This study models holobiome evolution using available gut microbiome information imputed from metadata from GenBank, the largest and most comprehensively annotated public genomic database from the United States National Library of Medicine. Based on clusterings derived from a phylogenetic similarity approach, we examined correlations between microbiome data with host diet and domestication. We also compared the oral and gastrointestinal (GI) microbiome across hosts. The results of this study demonstrate the potential to leverage available microbiome data to enable holobiome analyses.

## Results

The resulting presence-absence data matrix from GenBank had 184,659,278 entries, of which 37,007,067 were GI and 3,011,594 were oral host-microbe data. The GI matrix had 147 host taxa by 59 microbial phyla (consisting of 28,373 species) and the oral matrix had 51 host taxa by 25 microbial phyla (consisting of 4338 species). Seventy-six microorganism phyla were common across both datasets and 50 uncultured/unclassified phyla were excluded. Of the 163,905 unique microbial species, 9070 could not be identified and 885 uncultured specimens were removed. Analysis of GI dataset depicted Proteobacteria as the dominant phylum across omnivores and carnivores, with Firmicutes as the dominant phylum in herbivores, followed by Actinobacteria, Pisuviricota, Bacteroidetes, Apicomplexa, Ascomycota, and then others (Figure 1). Microbial composition from carnivores to herbivores shows a trend of decreasing Proteobacteria and increasing Firmicutes, as well as an increase in microbial diversity in herbivores.

**Figure 1.**
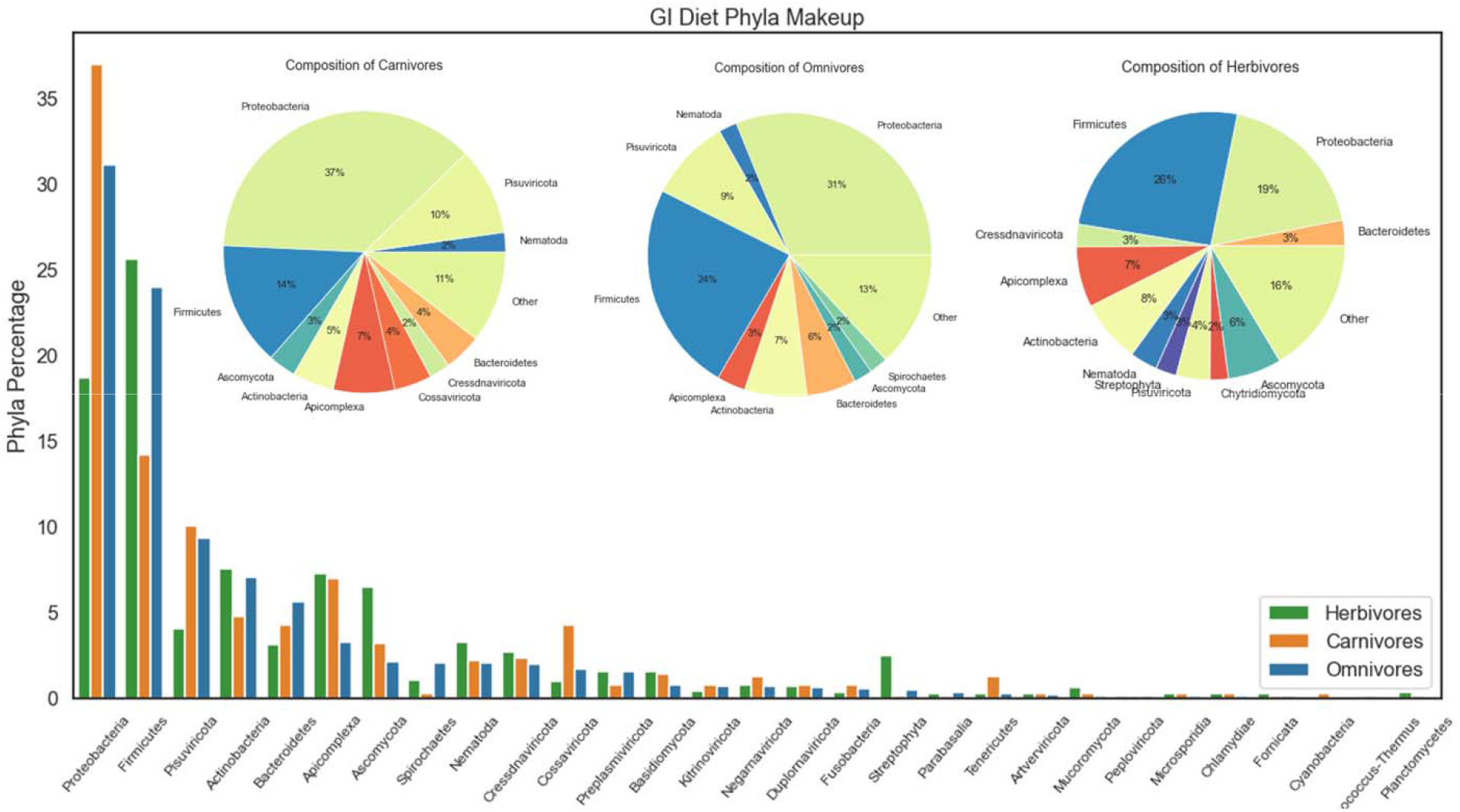
Comparison of individual microbial phyla (x-axis) found in gut samples. Phyla with at least one grouping above 0% were used. Pie charts of each dietary group’s composition is also shown, with any phyla under 2% representation merged into a general group, “Other.” An inverse relationship between Proteobacteria and Firmicutes is visible from carnivores to herbivores, with high Proteobacteria for carnivores and high Firmicutes in herbivores. Increased diversity is also visible in herbivore composition.

Across both GI and oral data, Proteobacteria were found in 96 of the 148 hosts (65%), followed by Firmicutes (53%) and Apicomplexa (52%). Certain clades show lower amounts of specific data, such as bats with only viruses and primates with mostly eukaryotic symbionts. In the oral microbiome data, Firmicutes show more dominance in herbivores and omnivores, while Proteobacteria leads the carnivore composition. The herbivores had more microbial diversity and Proteobacteria and Firmicutes followed an inverse relationship (Figure 2).

**Figure 2.**
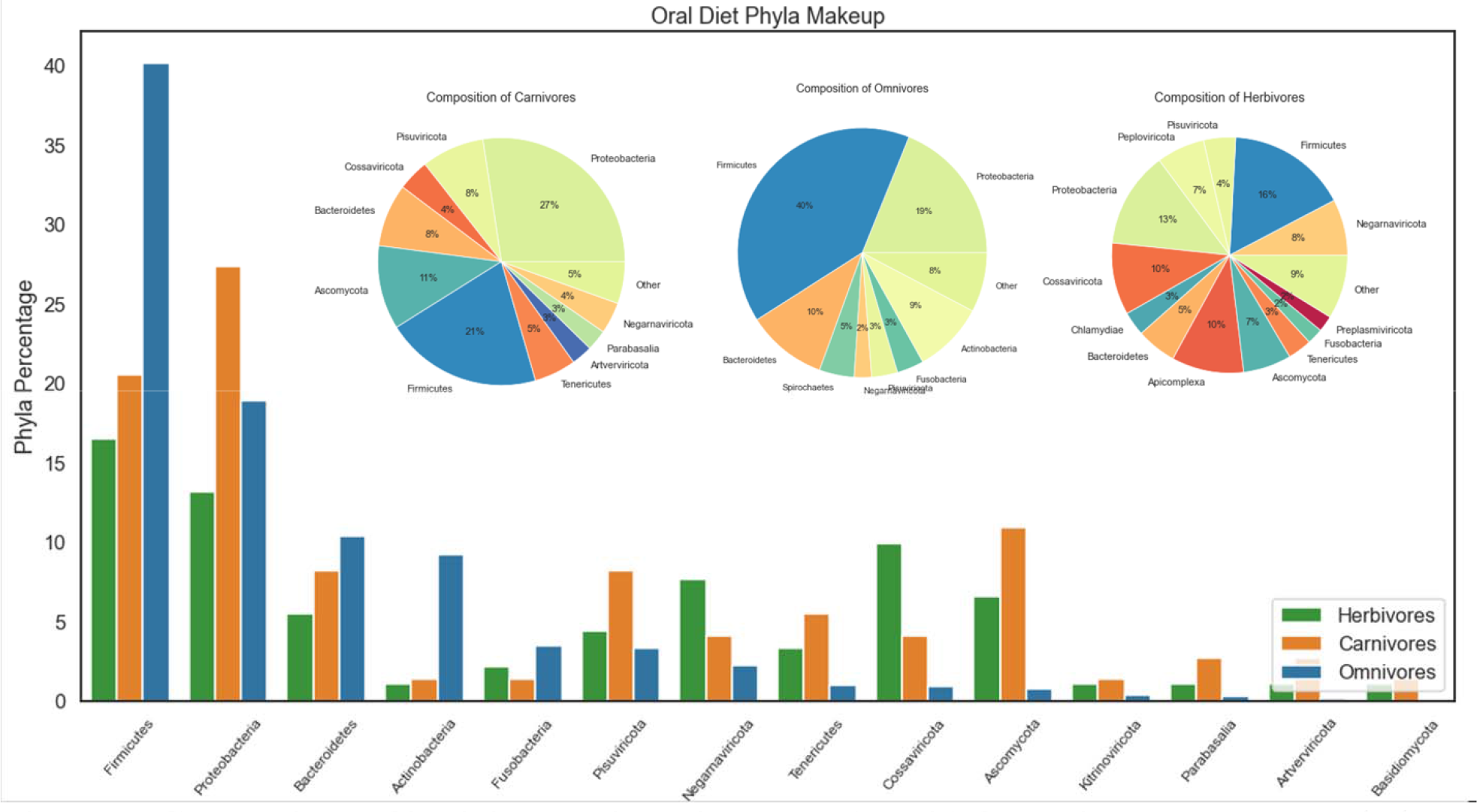
Comparison of individual microbial phyla (x-axis) found in oral samples. Phyla with at least one grouping above 0% were used and had much fewer phyla represented here compared to the gut data. Pie charts of each dietary group’s composition is shown as well, with any phyla under 2% representation merged into a general group, “Other.” Herbivore composition is visibly more diverse than the omnivore or carnivore microbial diversity. The same trend of inverse Proteobacteria-Firmicutes is not present here.

Based on the GI tree (Figure 3), humans are distinct from most primates, alongside domesticated omnivores including dogs, animal models, and livestock animals. Herbivorous mammals fell into two groupings, one with larger herbivores (e.g., buffalos and camels) and a mixed group (e.g., lemurs and deer), indicating more shared attributes. Carnivores and omnivores had no specific clustering. Ray-finned fishes (Actinopteri) were found near each other. Birds (Aves) were distributed across the clades. Rodents were also generally observed to be grouped together. Primates were mostly found together, noticeably missing humans and rhesus macaques. A tanglegram (Figure 4) analysis of the GI and oral holobiome trees included more omnivore and herbivore hosts than carnivores, and generally indicates a trend towards the consistent clustering of domesticated animals. This suggests that GI and oral microbiomes are more similar than different between the included domesticated animal hosts. Both sets of herbivores in the GI and oral trees showed the highest amount of microbial diversity across the three diet types and had more individual phyla. Only the omnivores exhibited a difference in which a given phylum dominated the oral and GI tract, where Firmicutes dominated oral and Proteobacteria dominated GI.

**Figure 3.**
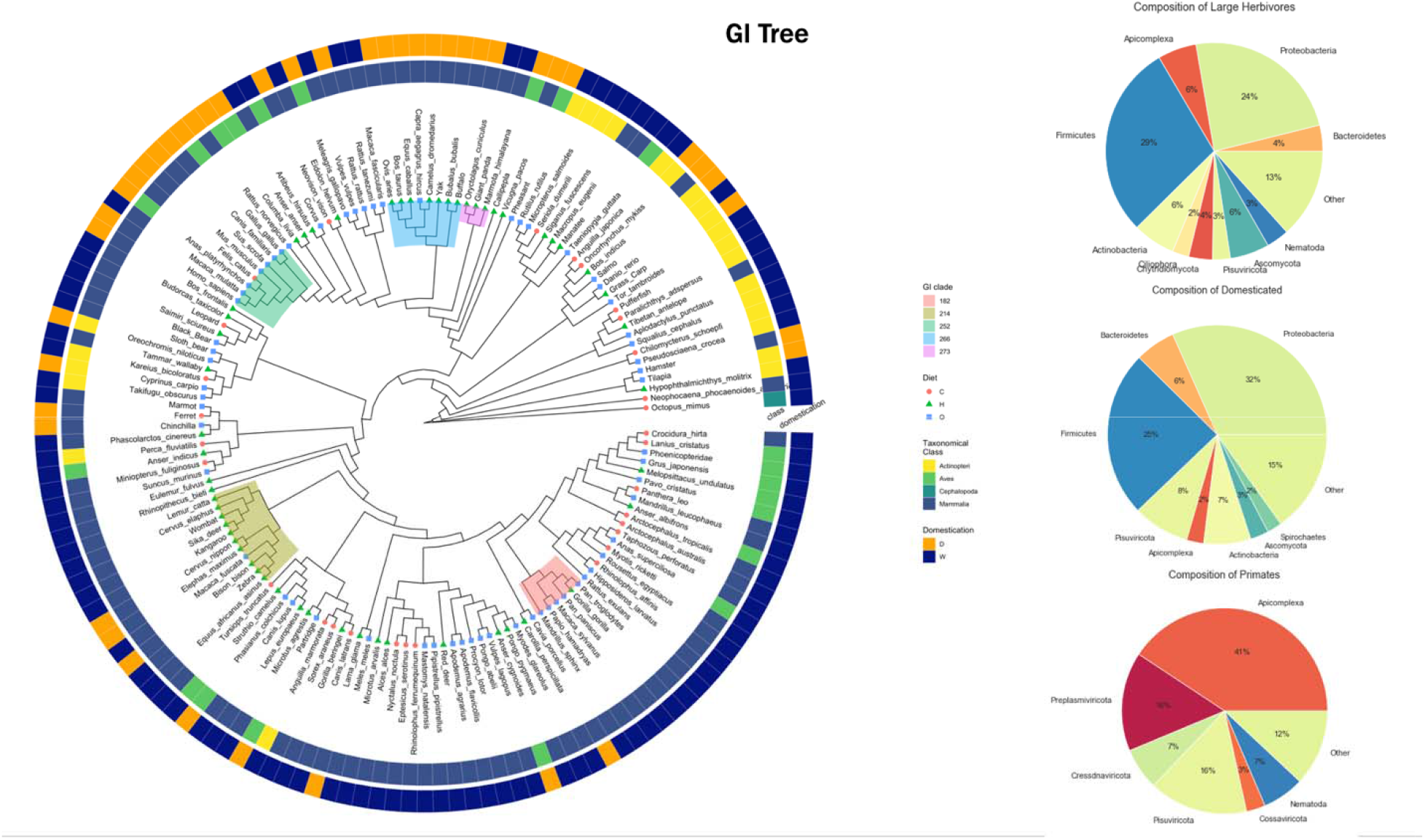
Phylogenetic analysis of the gut samples in host-microbiome data. Individual node are labeled with dietary grouping, with additional colored groups of taxonomy and domestication outside of the tree. Colored clades were chosen manually based on grouping by diet, domestication, and animal family. The herbivore, domesticated omnivore, and primates clade microbial composition is shown on the right, in which primate composition is primarily made up of eukaryotes and viruses, indicating a dataset bias.

**Figure 4.**
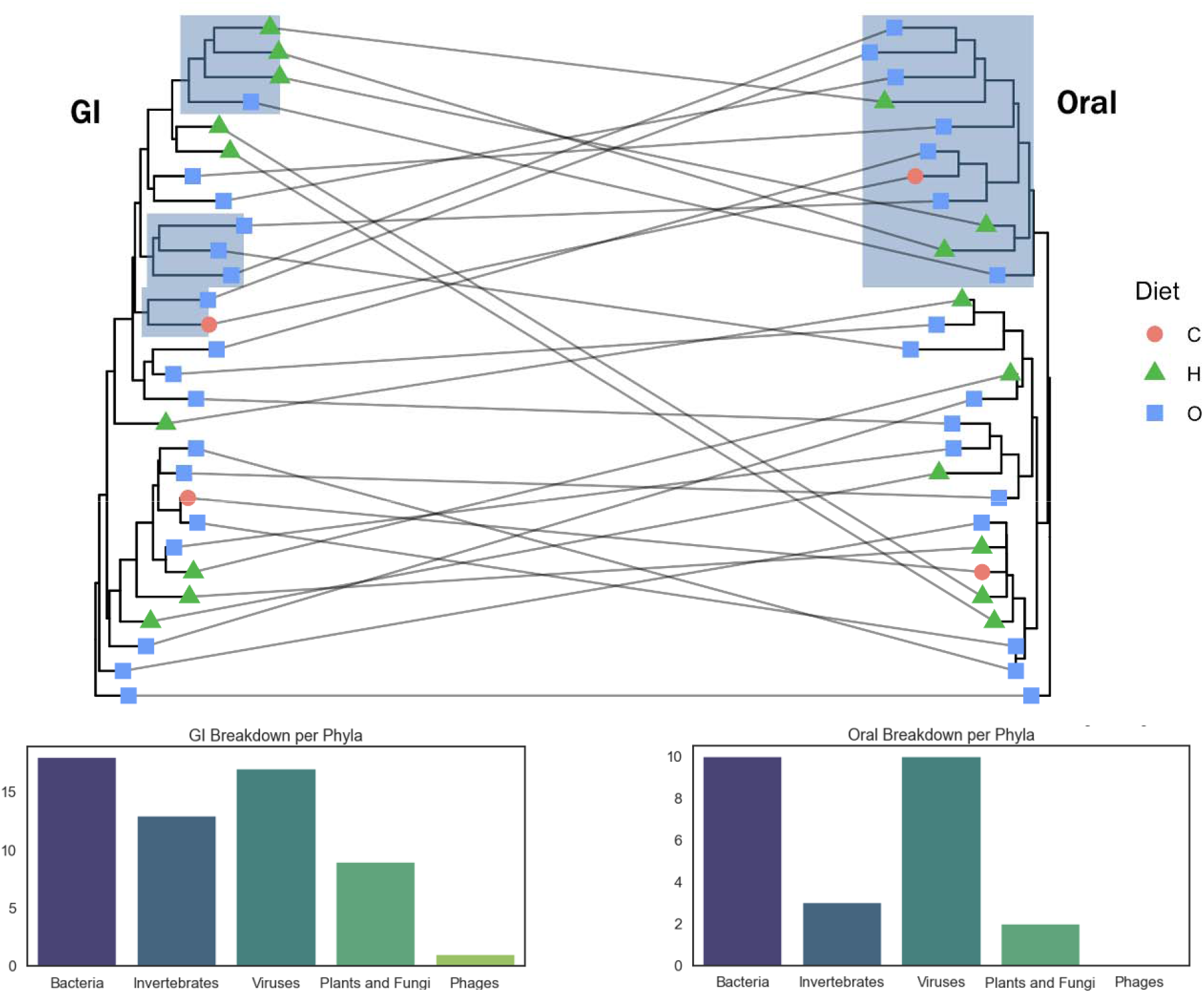
Phylogenetic analysis of the gut and oral samples in host-microbiome data. Individual nodes are labeled with dietary grouping and the highlighted clades represent the same domesticated grouping shared by the gut and oral trees. The separation of the clade in the GI tree demonstrates the split between herbivorous bovines and the other domesticated omnivores, whereas this group remains combined in the oral tree. Additionally, the breakdown of data per kingdom is shown for each dataset, in which Bacteria and Viruses dominate.

The three diet types indicated an inverse relationship of Proteobacteria and Firmicutes in both the GI and oral data. However, the omnivore group had a notable switch of Proteobacteria at the highest in the GI set and Firmicutes at the highest in the oral set. In contrast, both carnivores and herbivores had the Proteobacteria and Firmicutes dominate the GI and oral datasets, respectively.

## Discussion

This study used public gut microbiome data to model holobiome evolution. Our main findings support concepts previously reported and our core hypothesis that gut holobiome phylogeny correlates with diet. First, human microbiomes are distinct from other primates, as previously found [4, 14]. This distinction can be attributed to a gradual change in human versus other primate diets, including impact of the agricultural and industrial revolutions. Additional factors that could further explain the distinction of humans from other primates are the use of antibiotics, increased hygiene, and less exposure to wild microbes – all of which are implicated in the loss of microbial diversity. Second, dietary groupings were most observed in herbivores in the GI holobiome, which showed more consistency compared to non-domesticated omnivores and carnivores. The herbivore GI and oral holobiomes had the highest amount of microbial diversity, consistent with prior studies [15]. This is likely due to the myriad dietary fibers that strict herbivores eat, increasing microbial diversity distinct from omnivores and carnivores. Additionally, the lack of consistent grouping in our analysis of omnivores and carnivores is supported by previous studies, which indicate no significant differences between microbiome composition of these two diets [16]. Certain taxonomical groups showed fewer dietary markers: (1) Actinopteri tended to cluster at the bottom of the tree, although this could be due to limited data and (2) Aves were distributed across the tree, potentially due to varied omnivore diets. The dietary impact on the holobiome therefore appears to be most relevant to herbivores.

Our holobiome analysis method provides a unique outlook into how hosts are related by their microbiome data based on data available from GenBank. Phenetic approaches were chosen for this study because of the ambiguity of an absence datapoint. While a presence datapoint refers to the positive identification of the microorganism in a host, an absence datapoint is limited by what data are available in GenBank. Data limitations in specific hosts and the presence-absence data likely impacted the holobiome tree construction; however, current research supports that microbiome composition is not strictly segregated between different dietary groups [16]. An analysis by Youngblut, et al. similarly looked at the microbiome composition of hosts across animals using 16S RNA sequencing, revealing four distinct sub-networks based on presence-absence data: (1) Bacteroidetes, (2) Firmicutes, (3) Proteobacteria, and (4) Euryaarchaeota. Firmicutes and Proteobacteria were found in every species, followed by Actinobacteria and Bacteroidetes, as seen in our study. Proteobacteria were particularly dominant in carnivores, followed by higher levels of Bacteroidetes in omnivores and high levels of Firmicutes in herbivores. In ungulates and primates, Spirochaetes were identified as impactful, which were also found in our ungulate data but not primate data. Actinopteri were characterized by Proteobacteria, which was our data also confirmed. While Youngblut, et al. shows strong grouping by diet and taxonomic class, this is less clear with microbiome-based phylogeny and microbiome distinction across diets. The corroborating results between the Youngblut and our study demonstrate the promise in leveraging existing data repositories, such as GenBank.

The oral microbiome is a combination of environmental factors as well as digestive factors and has high microbial diversity, second only to the GI tract [17, 18]. The oral microbiome tends to have more Firmicutes and Fusobacteria, hosting more aerobic bacteria compared to the anaerobes in the gut [19]. Domesticated animals (e.g., dogs, cats, and other livestock) were observed to be consistent between the GI and oral holobiomes based on the tanglegram (Figure 5). Diet might change composition across host species, as it is known that oxygen and pH play the largest role in modulating bacterial colonization and biofilm formation in both the GI and oral microbiomes [20]. Perhaps due to standardized grains, increased hygiene, or similarity in processed foods, domesticated animals have greater microbiome similarity to one another compared to their ancestral species.

Previous studies on microbiome-host coevolution have shown that while the microbiome plays a role in shaping host evolution, the level of its impact can vary. Diet is known to be an important decider of microbiome composition [21]. Decreased quantities of dietary fiber, particularly seen in Western diets, are linked with more disease prevalence in humans. High-fat and high-sugar diets are associated with circadian rhythm disruption, which is also linked to the gut microbiome [22]. Gut microbiome functions of herbivorous and carnivorous mammals have shown differences in microbiome composition based on diet, in which herbivores tend to have more carbohydrate-metabolizing symbionts and carnivores have more protein-metabolizing enzymes [16]. Additionally, diet has been studied on an evolutionary scale. Studies have shown that while diet has little impact on diversification rates, there is relative conservation of diet across animals, perhaps due to the gut microbiome composition [23]. Thus, diet plays an important, yet ambiguous, role in holobiome evolution. Domestication can also impact gut microbiomes, yielding differences between wild and captive/domesticated animals [24]. Domestication impacts on animals have been paralleled with the impact of industrialization on humans, including changes in diet [25]. Plant microbiota also show reduced diversity under domestication, affecting crop production [26]. Because domestication is a nascent area of study with the gut microbiome, there are limited studies examining microbiome-host coevolution or impact of diet changes. Diet and domestication impacts can be paralleled by human microbiome research comparing vegan, vegetarian, omnivore, and high animal content diets [27]. Human diet research has shown that vegan and vegetarian diets are associated with more microbial genes for carbohydrate metabolism, amino acid synthesis, and short chain fatty acid and vitamin production [3]. In contrast, high animal protein diets show increased trimethylamine N-oxide and proteases [21]. This difference can be demonstrated in the gut microbiome genera composition of the two diets as well. High animal protein diets show increased bile-tolerant bacteria (*Proteobacteria, Bacteroidetes*) and plant protein diets have increased dietary plant polysaccharide fermenters (*Firmicutes*) [27]. *Bacteroidetes* and *Firmicutes* play roles in fiber digestion and short chain fatty acid production while *Proteobacteria* are commonly pathogenic but are also the most abundant in the human gut [28-31]. Given this difference in microbiome composition between human diets as well as herbivore and carnivore diets in the animal kingdom, diet and domestication can play a role in the holobiome.

In addition to supporting the development of a comprehensive holobiome database, our analysis identifies gaps and biases in the collection of microorganism data, especially in consideration of holobiome studies. The bacterial phylum Proteobacteria was the most represented phylum, which could be due to a bias towards clinical research since many Proteobacteria are pathogens. This bias is notable among the human microbiome data, where Proteobacteria and Firmicutes dominated. In contrast, Bacteroidetes and Firmicutes make up 90% of the abundance of the microbiome; however, this does not reflect the number of individual unique species present in each phyla and therefore it is unclear if our data composition is fully reflective of the human microbiome [32]. Additionally, Bacteroidetes has a smaller representation across our dataset. This phylum is well associated with omnivores and in humans, especially the genera of *Prevotella* and *Bacteroides*, where the former is seen with fiber heavy diets and the latter is characterized as an opportunistic pathogen in western diets [33]. It is possible that the decreased Bacteroidetes data is due to challenges in culturing anaerobic species. Additionally, diseased patients often have increased levels of Proteobacteria and decreased Bacteroidetes during analysis of their gut microbiomes [34]. Another area of low representation is Archaea. The phylum Euryarchaeota plays an important role in the gut microbiome and is normally characterized as one component of microbiome composition. However there is likely limited data on this phylum because Archaea are hard to culture and identify [32]. The remaining phyla are comprised of fungi, protozoa, viruses, and invertebrates, of which fungi and protozoa have been linked to digestive functions while viruses and invertebrates impact microbial diversity [35-40].

This study reveals the potential to harness publicly accessible data for advancing holobiome research. Our findings provide the basis for supporting holobiome evolutionary studies, as demonstrated here in the context of diets across the animal kingdom. Our results further support that the human holobiome is distinct amongst primates, evolving alongside domestication. The framework developed in this study is poised to enable subsequent holobiome studies.

## Materials and Methods

Microbiome and associated host information was imputed from GenBank metadata, which were systematically downloaded and extracted using a Java tool (genbank-loader [41]). The tool retrieved and extracted GenBank metadata into a MySQL database, which included a record of a given microorganism, along with its provided source site, the recorded host, and the tissue type. The data from the fields were mapped to standardized terms. Data processing was done in Python3 and Julia. Isolation source and tissue type data were mapped to standardized UMLS concepts using the MetaMap tool from the National Library of Medicine [42], merged into a single dataset including only isolation sources that were tissue types or organs, and then categorized into general groupings for tissue type source sites. The groupings of interest for this study were gastrointestinal (GI) concepts (feces, abdomen, intestine, rectum, stomach, esophagus, rumen, colon, ileum, cecum, jejunum, and duodenum), and oral concepts (mouth, saliva, tonsil, throat, enamel, tongue, and oropharynx).

Host data were processed using a combination of MetaMap and the NCBI Entrez Taxonomy database to map user entered data to scientific names of animal hosts. For this study, only hosts with gastrointestinal tracts were kept and later filtered to solely include the following classes: Mammalia, Aves, Reptilia, Amphibia, Cephalopoda, and Actinopterygii. Microbe organism data were mapped to NCBI Entrez Taxonomy database and used to retain bacteria, archaea, fungi, protozoa, viruses, and invertebrate parasites. Uncultured or unspecified bacteria and viruses of non-specific phyla were not included.

The combined data matrix was organized into a presence-absence matrix, where each host had a binary value indicating whether a particular phylum of microorganisms was found in the host data. Weight sampling was used to equalize coverage across all hosts. The probability distribution was calculated by measuring the number of records in the dataset for a microorganism per host and then used to randomly select data. Non-microbiota and previously defined host data were filtered out and all phyla without at least one datapoint were removed. GI and oral data were individually run through PAUP* (Phylogenetic Analysis Using Parsimony *and other methods) to create trees using the Neighbor Joining phenetic (distance-based) approach. Trees were calculated for both the oral and GI datasets, as well as a set of trees in which only shared hosts between both datasets were used. Trees were rooted to the host that had the earliest divergence from the rest of the hosts, based on phylogeny using TimeTree (http://timetree.org [43]).

The phylogenetic and tanglegram trees were visualized using ggtree and treeio in R, while integrating diet and domestication data to highlight clade groupings [44, 45]. Pie charts for groupings of herbivores, carnivores, omnivores, and other tree-specific clades were generated using Matplotlib Pyplot in Python3.

## Acknowledgements

The authors thank Matthew Storer for his technical assistance with the installation and use of the genbank-loader program. Additional technical support for this project was provided by the Center for Computation and Visualization at Brown University. This work was funded in part by funding from the National Institutes of Health (U54GM115677).

## Author Contributions

INS conceived of the study and provided guidance for all technical and analytic aspects of this study. SM and VR implemented the process for downloading GenBank data using genbank-loader, extracted the metadata of interest, and developed the pipeline for using MetaMap to standardize text strings to normalized forms, as well as generating the presence-absence matrix used by this study. VR performed the phylogenetic analyses and developed the graphics used for the interpretation of the results by INS and VR. VR led the drafting of the manuscript, with substantiative input from INS and SM.

